# Topological Excitations govern Ordering Kinetics in Endothelial Cell Layers

**DOI:** 10.1101/2024.09.26.615134

**Authors:** Iris Ruider, Kristian Thijssen, Daphné Raphaëlle Vannier, Valentina Paloschi, Alfredo Sciortino, Amin Doostmohammadi, Andreas R. Bausch

**Affiliations:** Lehrstuhl für Zellbiophysik E27, Technical University of Munich, 85748 Garching, Germany; Center for Functional Protein Assemblies (CPA), Technical University of Munich, 85748 Garching, Germany; Matter to Life Program, Max Planck School, München, Germany; Center for Organoid Systems and Tissue Engineering (COS), Technical University of Munich, 85748 Garching, Germany; The Niels Bohr Institute, University of Copenhagen, Blegdamsvej 17, DK-2100 Copenhagen O, Denmark; Department of Vascular and Endovascular Surgery, Klinikum rechts der Isar, Technical University of Munich, 80333 Munich, Germany; German Center for Cardiovascular Research DZHK, Partner Site Munich Heart Alliance, 80336 Berlin, Germany

**Author notes:** These authors contributed equally (Iris Ruider, Kristian Thijssen).

## Abstract

Many physiological processes, such as the shear flow alignment of endothelial cells in the vasculature, depend on the transition of cell layers between disordered and ordered phases. Here, we demonstrate that such a transition is driven by the non-monotonic evolution of nematic topological defects and the emergence of topological strings that bind the defects together, unveiling an intermediate phase of ordering kinetics in biological matter. We used time-resolved large-scale imaging and physical modeling to resolve the nature of the non-monotonic decrease in the number of defect pairs. The interaction of the intrinsic cell layer activity and the alignment field determines the occurrence of defect domains, which defines the nature of the transition. Defect pair annihilation is mediated by topological strings spanning multicellular scales within the cell layer. We propose that these long-range interactions in the intermediate ordering phase have significant implications for a wide range of biological phenomena in morphogenesis, tissue remodeling, and disease progression.

## Introduction

Since 1858, the relationship between a homogeneous endothelium and blood flow patterns has been recognized, with endothelial cells displaying high levels of alignment in the aorta, in contrast to their misalignment at bifurcation points [1, 2]. Subsequent *in vitro* studies revealed that endothelial cell layers have an intrinsic ability to realign and elongate in response to shear stress [3–5]. Since then, the extent of this tissue-wide global order has been considered a defining characteristic of various cell layers, influencing processes such as vascularization [6], tissue regeneration [7], development [8], and morphogenesis [9–11]. However, it is increasingly evident that disrupting the order in localized areas, known as topological defects, serves as a hotspot for biological activity such as the regulation of cell apoptosis [12], cell layer homeostasis, and stem cell accumulation [13–15]. In non-living systems, it is well-established that local defects are crucial for transitions between disordered and globally ordered phases [16]. Understanding those transitions has provided the basis for applications in material science, ranging from superfluidity and melting [17–19] to superconductors [20, 21]. Yet, transitions between local and global order in living, cellular systems remain largely unexplored, partly due to the large length and time scales involved [22]. Instead, our understanding remains limited to characterizing the steady-state properties of disordered or ordered cellular assemblies. This lack of knowledge about intermediary states between disordered and ordered states of cell organization leads to heterogeneous and sometimes contradictory observations, as seen in endothelial cell alignment [23–27]. To unravel the kinetics of shear flow-driven ordering in primary endothelial cells, we employed time-resolved large-scale imaging paired with physical modeling. We show that the flow-induced order transition in endothelial cell layers is governed by a three-stage process in which the ordering kinetic is disrupted by an intermediate stage where cells temporarily misalign, and multicellular defect strings emerge. We further demonstrate that an intricate interplay between the external shear flow and the activity of the living endothelial cell layer governs the emergence of defect strings and the concomitant intermediate ordering phase.

### Ordering kinetics characterized by three distinct phases

To gather sufficient statistical data to fully describe the phase ordering kinetics between disordered and ordered cellular organization, we examined the effect of a constant superjacent shear flow (≈20 dyn/cm^2^) on a confluent layer of human aortic endothelial cells (HAOEC) [5, 29, 30] with an automated microscope setup, which allowed stitching of high-resolution images on a scale of several millimeters during periods of up to 64 hours (Fig. 1a). During HAOEC alignment and elongation (Ext. Data Fig. 1), the emergence of nematic order was observed in phase contrast images (Fig. 1b,c and Supplementary Movie 1). To quantify the order, we extracted the local nematic director field of the endothelial cell monolayer in the full field of view (Fig. 1b,f). From this, we obtained the orientation angle α, color-mapped throughout the field of view to spatially resolve and visualize the alignment. We identified the presence of nematic +1/2 and −1/2 defects within the cellular monolayer (Fig. 1c,f,g,h). Upon application of shear flow, an initially disordered layer of cells evolved into a well-aligned ordered layer, and the density of topological defects dropped significantly from over 400 defect pairs in the initial state to around 0 at the final state. (Fig. 1i). While in passive materials, the ordering kinetics would be characterized by a monotonous decrease of the topological defect density [31], here we observed three distinct phases driven by the non-equilibrium nature of the cell layer: In the initial ordering phase (I) exposure to shear stress induced the cells’ elongation and alignment along the flow direction. This was accompanied by a decrease in defect density in the first nine hours after flow was applied. In the subsequent intermediate disordering phase (II), from t = 10 and t = 25 hours, a transient reduction in cellular alignment and an increase in defect density was observed. In the final ordering phase, which we call the steady-state ordering phase (III), the system realigned again, almost all nematic defects disappeared, and a long-range orientational order was established (Fig. 1i and Supplementary Movie 1).

**FIG. 1.**
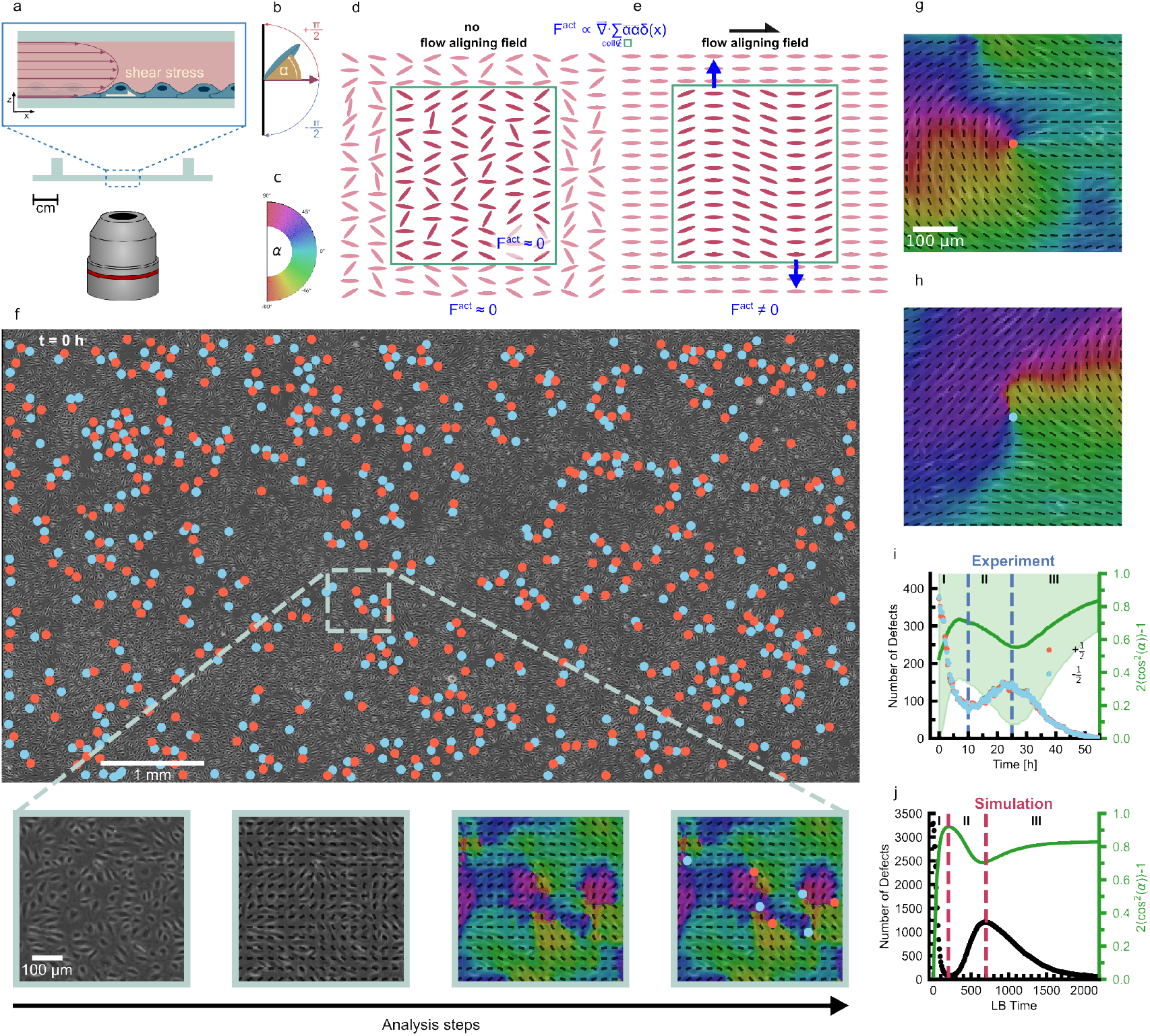
Shear flow induced alignment and defect dynamics in endothelial cell monolayers. (a) The cells experience a shear flow along the x-z plane. (b) Cell alignment is captured by the nematic director field, where the orientation α = 0 is defined with respect to the direction of the shear flow. (c) The nematic director is color-coded to visually represent cell alignment. (d) Without a flow-aligning field, inter-cellular forces are randomly oriented and cancel each other out. (e) The shear in the x-z plane acts as an affine alignment field, inducing elongation and ordering of the cells along that direction. As local ordering is established, inter-cellular forces stop canceling out, resulting in mesoscopic active forces F ^*act*^ (collective active stresses), approximated as extensile forces (ζ > 0) [28]. (f) Field of view of the endothelial cell monolayer during a flow experiment. The zoom-ins show a phase contrast image of a cellular monolayer and simultaneously illustrate analysis steps with increasing details containing the nematic director field, the orientation field, and nematic topological defects. (g) A representative +1/2 defect indicated by a red dot. (h) A representative −1/2 defect indicated by a blue dot. (i) The number of defects and the global nematic ordering of the endothelial cell monolayer during the alignment process in the experiments. The light green area indicates the standard deviation from the mean. (j) The number of defects (black dots) and the global nematic ordering of the endothelial cell monolayer during the alignment process in the simulations.

### Interplay between local activity and global shear

The appearance of the intermediate disordering phase suggests that while shear flow drives the ordering of the cells, the active motion and cell-cell interactions within the endothelial layer oppose the induced ordering. To test this, we simulated the transient behavior using active nematic theory (‘Simulations’ in Materials and Methods). Here, we used a coarse-grained model to solve the local velocity 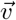 and the nematic orientation tensor **Q**. The superjacent shear flow was modeled as an external aligning field that energetically penalizes orientations perpendicular to the flow direction, similar to an anisotropic quenching field [32]. This induced a global nematic orientation of the cell layer. We modeled the activity of the endothelial cells by out-of-equilibrium force contributions, often called active force, dominated by dipolar forces along the resulting local elongation axis (Fig. 1d,e). The activity and anisotropic field competed at different timescales, which resulted in a variety of behaviors (Fig. 2), including the non-monotonic coarsening behavior (Fig. 1j). As the cells elongated under the shear flow and local orientational order was established, the active stress from fluctuations in the orientation field was sufficient to nucleate topological defects and transiently destroyed the local orientational order in the system. Eventually, the orientation of the cells became almost perfectly aligned along the preferred axis set by the external field.

**FIG. 2.**
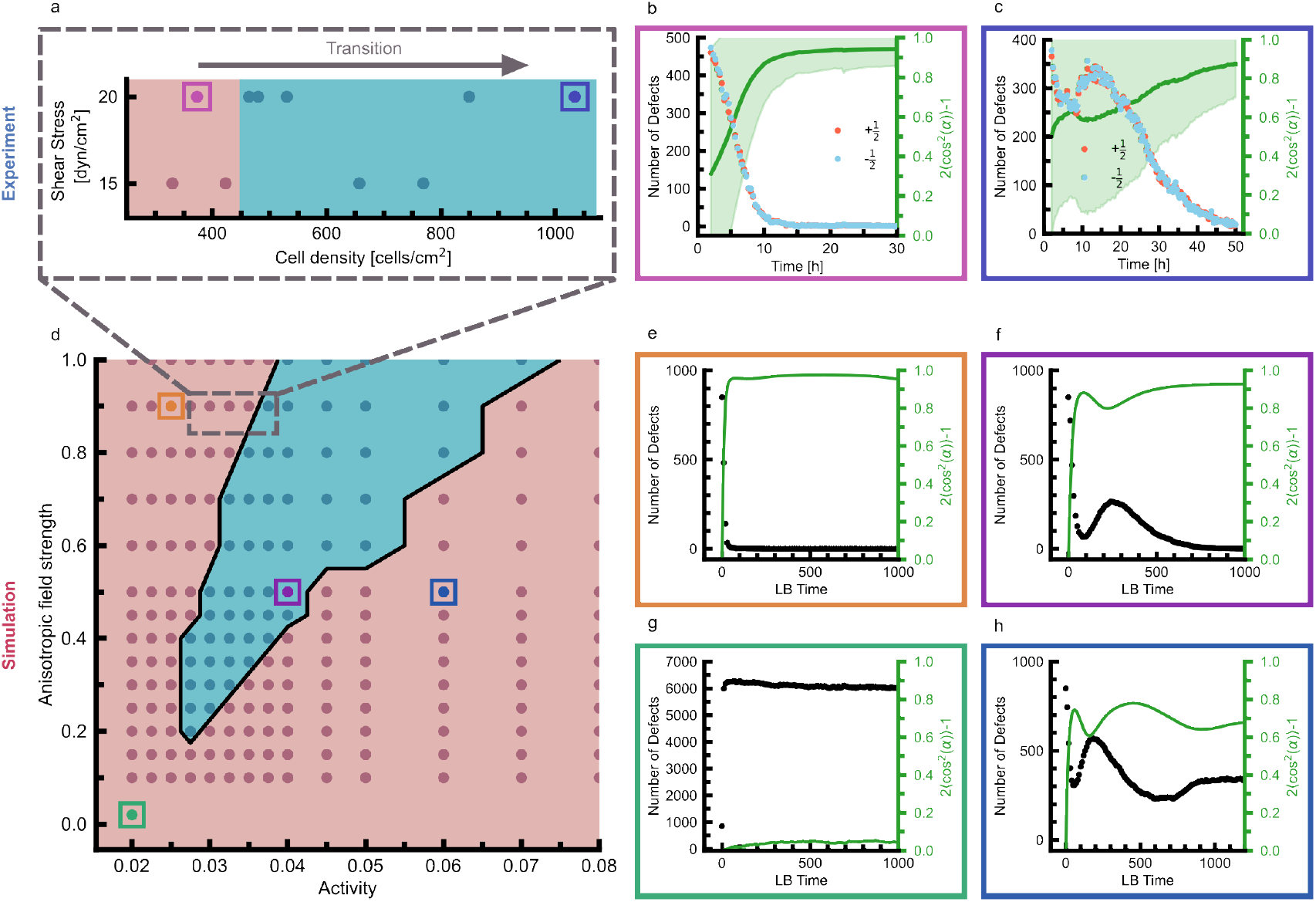
Interplay between cellular activity and external aligning field governs alignment dynamics. (a) The phase diagram for the experimental data obtained from several experiments at different cell densities shows a transition from the regime with monotonous alignment dynamics in red towards the regime with a non-monotonous alignment behavior in blue. (b) At lower cell densities, the endothelial cells align monotonously along the flow direction, lacking the three-stage dynamics. The light green area indicates the standard deviation from the mean. (c) At higher cell densities, the endothelial monolayer displays non-trivial alignment dynamics characterized by a non-monotonous alignment. The light green area indicates the standard deviation from the mean. (d) The phase diagram obtained from simulations shows that the non-trivial regime, highlighted in blue, only exists within a very defined region. The trivial region is highlighted in red. (e) The system displays a monotonous alignment behavior in the trivial region for high anisotropic field strength and low activity within the nematic director field. (f) In the non-trivial regime, the interplay between the anisotropic field strength and the intrinsic activity gives rise to a threestage, non-monotonous alignment behavior. (g) When activity and flow anisotropic field are both low, the cells don’t display localized order, resulting in an isotropic state. The number of defects saturates as orientational order becomes ill-defined. (h) When the activity is the dominating parameter, the system fails to align and evolves towards a state with only local nematic alignment.

In our experimental setup, rising cell density entailed an increase in the activity of the monolayer due to enhanced cell migration and cell-cell interactions, which collectively augmented the intrinsic active stress [33], if cell density was low enough that no jamming occured. Experiments carried out at different cell densities revealed that at low cell densities and thus lower active stress, endothelial cells aligned monotonously in the flow direction as the external field dominated the system (Fig. 2 a,b). However, with increasing cell density and thus increasing activity within the cellular monolayer, we observed the emergence of an intermediate phase during cell alignment (Fig. 2 a,c). While the experimental setup offered only a limited range to explore this transition, simulations provided a more comprehensive understanding of how the activity of the monolayer and the strength of the external flow field influence the nature of the cell ordering process. The phase diagram illustrates how the interplay between activity and the external field’s strength affects the kinetics of transition between disordered and ordered states. In congruence with experiments, and as expected for passive materials, in the absence of or at low values of active stresses, when the aligning field dominates on all time scales, the ordering kinetic follows a monotonic rarefaction of topological defects (Fig. 2d,e). In contrast, larger active stresses result in a defect-loaded steady-state, called active turbulence (Fig. 2d,h). Only when the active stress and the external field compete on similar timescales can we recover the experimental behavior, where the global nematic order parameter exhibits three distinct transient phases (Fig. 2d,f). We point out that for a weak anisotropic aligning field, the system remains in an isotropic state with no emergence of global order (Fig. 2g). The existence of a threshold shear stress, below which endothelial cells no longer respond to flow, has already been reported in the literature [26, 34]. Therefore, only in an intermediate regime does the intricate interplay between activity-driven defect nucleation and flow alignment forces result in an intermediate increase in defect numbers.

### Anisotropic correlation lenghts

Because of the preferred orientation axis set by the direction of the superjacent flow, the coarsening of the local nematic order was not isotropic. Rather, the system can be split into domains of negative and positive rejection to the preferred axis δn, which divides these domains (Fig. 3 a,e). The multicellular orientationally ordered domains slowly coarsened in an anisotropic manner as global order was established (Supplementary Movie 2 and 3). This is best quantified by measuring the correlation functions 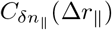 and 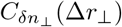 that resolve the orientation correlation along the perpendicular Δr_⊥_ and parallel Δr∥directions to the preferred axis (‘Correlation Functions’ in Materials and Methods). The correlation lengths found from the correlation function along the parallel ξ∥and perpendicular axes ξ_⊥_ to the preferred orientation showed asymmetric growth of similarly aligned regions in both the model and experiments, where coarsening of the correlation length occurred primarily along the preferred axis (Ext. Data Fig. 2a,e). Interestingly, the domain growth along the axis parallel the flow direction was not uniform in time and was closely linked to the three previously observed ordering phases, as we discuss next.

**FIG. 3.**
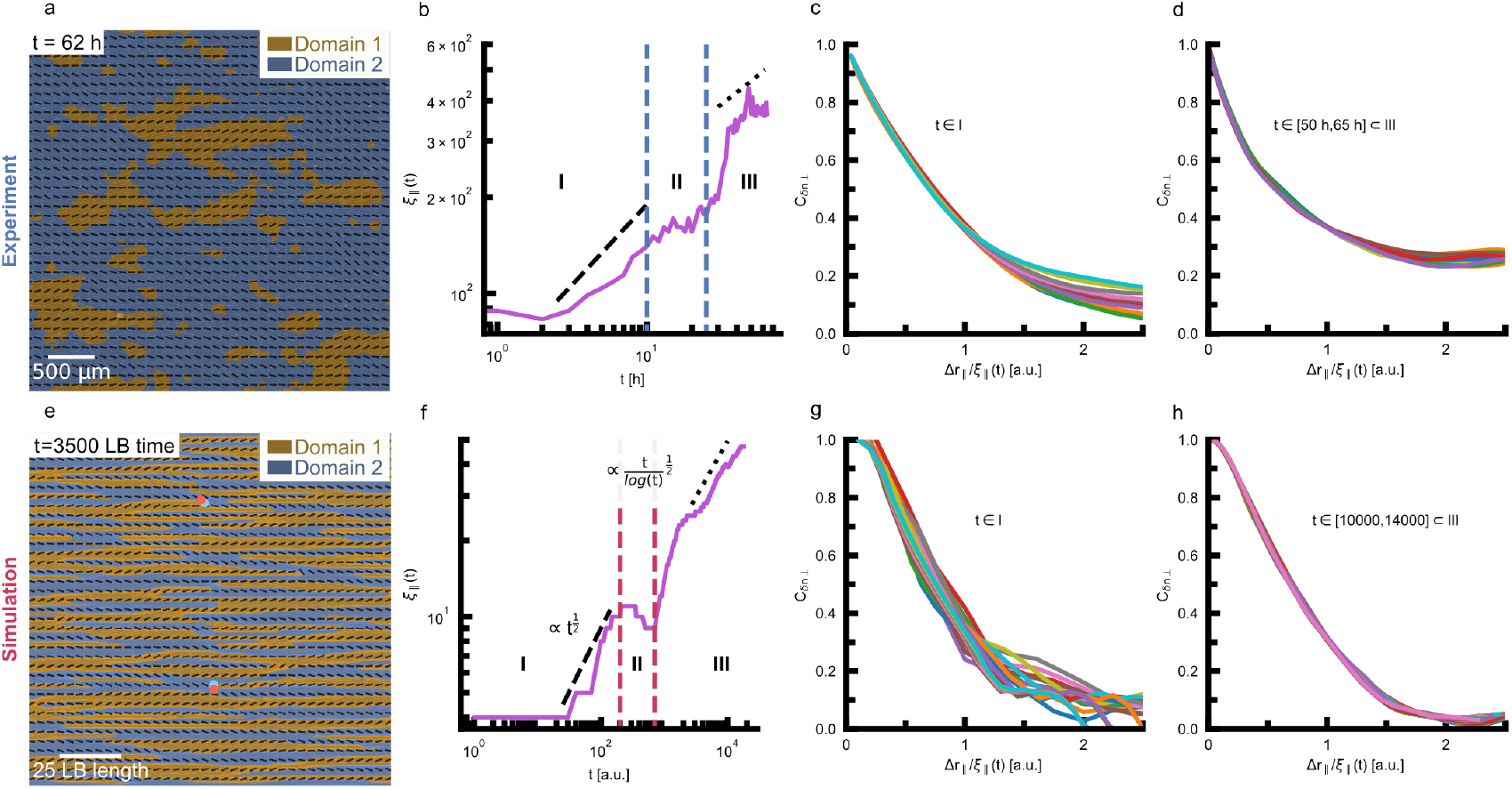
Three distinct scaling behaviors of the domain growth. (a) Illustration of the domains with positive or negative projection 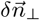 values corresponding to the experimental data extending anisotropically along the flow axis. (b) The correlation length ξ_∥_ exhibits distinct growth rates. (c) The correlation functions normalized by the length scale ξ_∥_ during the initial ordering phase (I) for the experimental data. (d) The correlation functions normalized by the length scale ξ_∥_ for late times during the steady-state ordering phase (III) for the experimental data. (e-h) Corresponding simulation figures to (a-d). In (b) and (f) the dashed and dotted lines denote 1/2-power law scaling and 1/2-power law scaling with logarithmic correction, respectively.

### Three-stage kinetic ordering

In the initial ordering phase (I), orientation correlation functions along the preferred axis r_∥_ decayed to zero, and the characteristic correlation lengths ξ_∥_ increased exponentially. From the simple dimensional analysis of diffusive isotropic-nematic transitions [35], the growth of ξ∥ is expected to follow t^1*/*2^ which is consistent with our simulation results and close to the experimental data (Fig. 3b,f). This suggests a negligible impact from defect annihilations on the ordering kinetic in this initial phase. As the correlation functions did not collapse with the distance normalized by the correlation length Δr∥ /ξ∥, the growth of the ordering did not exhibit dynamic scaling behavior, that would be expected for kinetic ordering in passive materials (Fig. 3c,g) [36].

In the subsequent intermediate disordering phase (II), the asymptotic value of the correlation function at long distances did not drop to zero. It reached a finite value that increased with time, indicating the appearance of a true long-range order (Ext. Data Fig. 2e,h). The characteristic correlation length ξ∥ ceased to increase and remained stable while the number of defects rose, which is attributed to the increase of cellular forces as cells begin to align. As the active forces reached a critical value, an active instability set in which caused defects to nucleate [37].

In the steady-state ordering phase (III), the correlation functions converged to a finite value and entirely collapsed with Δr∥ /ξ∥, indicating a dynamical scaling of length-scales as the system became more ordered (Fig. 3d,h). The characteristic correlation length ξ∥ increased again, and for final times, this length scale is expected to follow 1/2 power law with a logarithmic correction due to the interaction between topological defects [31, 36]. While, due to the limited experimental accessible timespan, this precise form of temporal scaling could not be unambiguously determined in the experimental data set, it was possible to use prolonged simulations to confirm the existence of a logarithmic correction to the 1/2 power law growth in the final ordering phase (Fig. 3b,f). This indicated that the coarsening in the final stage was dominated by long-range logarithmic interaction between defects [36].

### Heterogenous distribution of bound defect pairs

The emergence of three-stage ordering kinetics, linked to the non-monotonic evolution of topological defects during the establishment of long-range orientational order (Fig. 1i,j, and Fig. 3b,f), prompted us to investigate the spatiotemporal defect dynamics leveraging the extensive field of view. A detailed examination of defect dynamics revealed that the emergence of new defect pairs was highly localized at the onset of the defect nucleation phase. During this phase, multiple defect pairs nucleated in close proximity to one another, while a significant proportion of the field of view remained defect-free. We characterized this heterogeneity by analyzing the distance between the k^th^ nearest neighbors of oppositely charged defects (Fig. 4a). We observed both in experiment and simulation that after defect pairs nucleated, they did not move further apart as expected in traditional active turbulence, nor did they immediately annihilate, as seen previously in flow-tumbling active systems with external fields [32]. Instead, the defects remained relatively bound to each other (Fig. 4b,c; k_0_). Hence, in contrast to a homogeneous increase in defect spacing, we identified regions densely populated with defects interspersed with void areas where no defects were present (Fig. 4d). This heterogeneity in the spatial defect distribution impacted the later stages of ordering kinetics. Consequently, during the final coarsening, the distance between nearest neighbor defect pairs k_0_ did not increase. Instead, the distance of the higher-order pairs k_*i>*0_ increased as the distance between defect-rich regions increased (Fig. 4b,c; k_*i>*0_). Eventually, at late times, the logarithmic interactions between the separate defect pair configurations became dominant, and all topological defect pairs annihilated. Even as the number of defects decreased and the domains grew larger (Ext. Data Fig. 3), we did not observe a significant change in the nearest neighbor distance between oppositely charged defects. This is because defect pairs tend to cluster in groups, and only these groups become increasingly sparse. This unique behavior did not occur in the absence of activity or the external flow field, indicating an intricate interplay between the cell activity, aligning field, domain interfaces, and the topological defects residing on them.

**FIG. 4.**
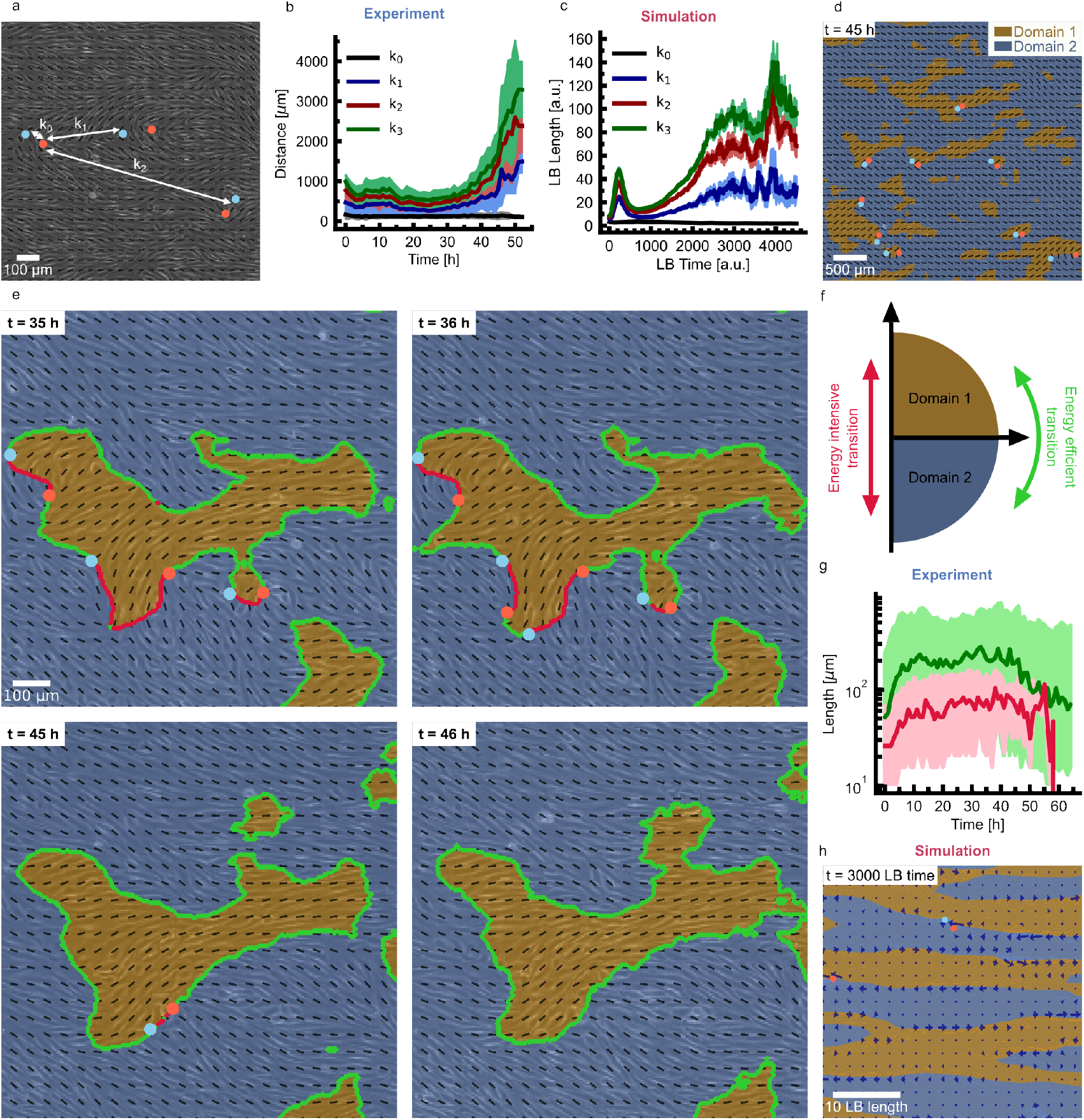
Nematic defects interact through strings. (a) The distance to the k^th^ nearest neighbour between two defects of opposite charge. (b) The time evolution of the distance between the k^th^ nearest neighbors for the experimental data. The brighter areas indicate the standard deviation from the mean for the corresponding color. (c) The time evolution of the distance between the k^th^ nearest neighbors for the simulation data. The brighter areas indicate the standard deviation from the mean for the corresponding color. (d) In the experimental data, the nematic defects sit on the domain boundaries. The nearest neighbor pairs remain at a close distance from each other, while the distance to the second nearest neighbor increases during coarsening. (e) The nematic defect sitting on the domain walls interacts through strings with parallel orientation (green) and perpendicular orientation (red) to the preferred orientation axis. (f) The transition from one domain to the other can be energy intensive, indicated by the red arrow, or energy efficient, indicated by the green arrow. (g) The average length of the strings with perpendicular orientation (red) decreases to zero as the system coarsens. The length of the green string initially increases and eventually plateaus during coarsening for experimental data. The brighter areas illustrate the interquartile range for data of the corresponding color. (h) Force field of the nematic director field. The forces (blue arrows) are highest along the domain boundaries.

### String excitations

In the absence of an external field, it is well-established that activity unbinds pairs of oppositely charged nematic topological defects [37, 38]. Interestingly, in both experiments and simulations, we measured a fixed, small distance (≈123 *±*11 µm, equivalent to up to 11 cell sizes) between nearest neighbor k_0_ oppositely charged *±*1/2 defects throughout the ordering process. This indicates that, despite the intrinsic activity of the cells, the oppositely-charged defect pairs remained bound during coarsening, even after the nucleation phase. We hypothesized that the emergence of such stable defect configurations is governed by topological strings that bind defect pairs together [39], which are driven by the interplay of active stresses within the cell layer and the external aligning field.

To test our hypothesis, we identified regions that separated orientation quadrant 1 (domain 1, with positive δn_⊥_) from quadrant 4 (domain 2, with negative δn_⊥_) (Fig. 4e,f). Along these domain boundaries, the director angle α displayed maximum or minimum gradients in reference to the preferred axis. Remarkably, in both experiments and simulations, we found distinct strings between oppositely charged topological defect pairs, binding them together. Due to the symmetry-breaking field, we could define strings as director isolines that connected two oppositely charged defects along the preferred direction. In general, in the absence of an external flow, an infinite number of isolines exist along all possible directions, all of them equivalent. However, with the presence of a preferred direction, the isoline along the preferred direction gains physical significance as there is a different energy cost associated with the director transitioning from domain 1 to domain 4 through the preferred axis (green lines) compared to perpendicular transitions (red lines) (Fig. 4f). The evidence for this physical significance is manifested by the localization of the topological defect trajectories on these isolines in the experiments (Fig. 4e). As a result, string excitations stabilized the oppositely-charged defect pair configurations until the final ordering phase. Consequently, since defect motility was highly localized to movement along the strings, the strings provided a path for the annihilation of defects (Supplementary Movie 4 and 5), and the strings that lined up perpendicular to the preferred axis (Fig. 4g; red) eventually decayed as all defects were annihilated at later times. Meanwhile, the strings extending along the preferred axis (Fig. 4g; green) increased during the course of ordering, but the average string length appeared to remain constant, as small strings continuously nucleated. Examining snapshots of the ordering kinetics, however, revealed that eventually, long strings extending along the preferred axis dominated the system at late times (Supplementary Movie 4 and 5). The large extent of these strings is remarkable as they potentially provide a path for long-range transmission of mechanical forces. While force measurements were not available for the experiments, we could probe this mechanism in the simulations. Indeed, simulations confirmed that the strings act as hotspots for focusing active forces (Fig. 4h).

In summary, the high temporal and spatial resolution achieved in our experimental setup allowed us to describe the ordering kinetics of endothelial cell layers within the framework of active nematic liquid crystals, highlighting the governing role of topological excitation and explaining endothelial cell alignment as a dynamic transition with cellular misalignment as an intermediate stage. The introduced unifying framework reconciles previous contradictory observations reporting perpendicular or parallel endothelial cell alignment depending on experimental parameters [25– 27, 40–42]. The importance of the transient regime in endothelial cell alignment reveals the need for precise temporal resolution in detecting changes in protein expression, exemplified by the previously observed transient upregulation of JNK2 in bovine aortic endothelial cells [43]. The identified interplay of the activity-driven local nucleation of defects and global ordering induced by the external driving field sets the spatiotemporal behavior of the endothelial cell layers, which defines a phase diagram with distinct regions of tissue responses. This interplay gives rise to defects that are not fully localized rather they are connected by string excitation in endothelial layers and are the biological analogs of the pair-superfluid phase predicted for two-dimensional antiferromagnetic condensates under a magnetic field [44]. In this context, string excitations have been identified in classic XY models of magnetism, which possess both nematic and polar interactions [39, 44, 45] and are characterized by a linear interaction potential, in addition to the normal logarithmic Coloumb potential between the defects. As such, the linear potential leads to an effective line tension proportional to the length of the string binding the defect pairs together. The emergence of topological excitations spanning multiple cell lengths during the ordering process underscores the need for precise, localized measurements of protein expression and cellular responses at both local and global scales over time. The recent identification of a nematic phase during embryogenesis across multiple species, along with the observed presence of nematic defects during this process, highlights the potential significance of topological defects in development [46]. Our findings may set a benchmark for a wide range of biological phenomena in morphogenesis, tissue remodeling, and disease progression, where spatiotemporally localized mechanical effects have consequences over long distances and time scales.

## ACKNOWLEDGEMENTS

We gratefully acknowledge financial support by the European Research Council (ERC) under the European Union’s Horizon 2020 research and innovation programme (grant agreement no. 810104-PoInt). This research was conducted within the Max Planck School Matter to Life supported by the German Federal Ministry of Education and Research in collaboration with the Max Planck Society. This work was further supported by the Novo Nordisk Foundation grant no. NNF18SA0035142, NERD grant no. NNF21OC0068687 (AD), the Villum Fonden Grant no. 29476 (AD), and the European Union via the ERC-Starting Grant PhysCoMeT (AD) and the Horizon 2020 research and innovation programme Marie Sklodowska-Curie grant no. 101029079-SIMMS (KT).

## AUTHOR CONTRIBUTIONS

I.R. performed experiments with the contributions of D.R.V. and the support of V.P. under the supervision of A.R.B. K.T. and performed the simulations under the supervision of A.D. A.R.B., and A.D. conceived the project. Data Analysis was performed by I.R. and K.T. with the support of A.S. All authors participated in the writing of the manuscript.

## COMPETING INTERESTS

The authors declare no competing interests.

## I. MATERIALS AND METHODS

### Primary human aortic endothelial cell culture

Primary human aortic endothelial cells (HAOEC) (PeloBiotech, PB-CH-180-2011) were cultured in gelatine-coated T75 flasks (0.2 % gelatine in PBS incubated for 1h at 37 *°*C) using Endothelial Cell Growth Medium Kit Classic (PeloBiotech, PB-MH-100-2190) from passage 5 up to passage 9. For shear flow experiments, µ-slide I Luer ibiTreat (Channel height: 0.2mm) were incubated for 1h with 40µg/ml collagen at 37 *°*C. After three washing steps with PBS, 50 µl of HAOECs at a seeding concentration of 3 *·* 10^6^ up to 6 *·* 10^6^ cells/ml were seeded into the flow channel and left at 37 *°*C for 1 h to allow them to form first adhesions. Then, the cell culture medium was added to the reservoirs, and the HAOECs were left overnight to adhere fully to the substrate. The confluent cell layer resulted in 12 000 up to 40 000 cells per a field of view of 40 mm^2^. The cell culture medium, PBS, and the flow channel slides were kept in the incubator one day before cell seeding to prevent bubble formation in the flow channels.

### Perfusion cell culture

To avoid bubble formation, the cell culture medium and perfusion set were kept in the incubator one day before the flow experiment. The ibidi pump system and perfusion set (ibidi: orange) were prepared according to the manufacturer’s instructions. After the pump system was calibrated, the flow channel was connected. To allow the cells to adapt to the shear stress, a flow program slowly increased the flow velocity until reaching the desired shear stress of 15 dyn/cm^2^, respectively 20 dyn/cm^2^ after 1 h.

### Live cell imaging

Phase contrast imaging was performed using a Leica DMi8 Thunder Imager and a 10x (0.32 NA) air objective (Leica). To gather sufficient statistical data, we stitched high-resolution images over a scale of several millimeters for up to 64 hours. The area of the resulting field of view was 40 mm^2^. The ibidi stage top incubation system was adapted for live cell imaging with a custom-built lid to accommodate the tubing. The fluidic unit of the pump system was kept in a nearby incubator at 37 *°*C and 5 % CO_2_. Through a small opening in the rear of the incubator, the flow channel was mounted on the microscope stage while remaining connected to the pump system. Images were acquired every 20 minutes over at least 2.5 days.

*×*

### Characterization of nematic director field

We extracted the nematic field from the phase contrast images using a custom Python3 script based on the method from reference [47] as previously described in [48]. We computed the local nematic director ***n*** = ***t*** ⊗ ***t*** using the tangent vector ***t*** = (I_*y*_, −I_*x*_) to the gradient of the intensity of the image ∇ I(x, y) = (I_*x*_, I_*y*_) within a box of dimensions L *×* L. From the local nematic director, we extracted the eigenvalues n_1_ > n_2_ and the corresponding eigenvector ***e***_1_ to the biggest eigenvalue, which provided the nematic director associated with the pixel in the center of the box. Here, we used a box of 130 µm × 130 µm to compute the nematic director for a pixel. The nematic director field was computed for each pixel, and the spacing between two pixels is ≈ 5.2 µm.

### Detection of nematic defects

We obtained the **Q**tensor from the nematic director field. The defect cores were then identified as the local maxima, respectively minima, of the charge density 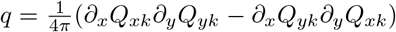 where |q | > 0.1. Only every 5^th^ pixel in the nematic director field was considered for defect detection.

### Extraction of nearest neighbour distance

Defects were considered for k^th^ nearest neighbor (knn) analysis if the defects were present for at least 4 h. We used trackpy to obtain and filter the defect trajectories.[49] From the defect trajectories, we constructed the knn-distance matrix, which we used to extract the knn distance for k = 0, 1, 2, 3.

### String visualization in experiments

The nematic director n = [cos(α), sin(α)] was defined both in orientation region region 0 ≤ α_1_ ≤ π (first and second quadrant) and −π/2 ≤ α_2_ ≤ π/2 (first and fourth quadrant). The parallel and perpendicular strings were defined to go between two resolution points if the absolute angle difference |Δα_1_| or |Δα_2_ | was larger than π/2, respectively.

### Length measurement for parallel and orthogonal strings

From the absolute angle difference obtained for string visualization, a binary image was created by setting the points with |Δα_1_| or |Δα_2_| larger than π/2 to one and any other point to zero. Eventually, string length was measured by obtaining the skeletons from the segmented features.

### Correlation functions

Taking the projection of the orientation parallel to the preferred axis δn (Ext. Data Fig. 2a) allowed us to identify how far away the director was from the preferred axis. Using this, we observed how regions of similar orientation coarsened by looking at the correlation functions Cδn_⊥_(Δr_∥_) (Ext. Data Fig. 2b) and Cδn_⊥_(Δr_⊥_) (Ext. Data Fig. 2c), where we split up the correlations along the perpendicular Δr_⊥_ and parallel Δr∥ direction to the preferred axis.

Comparing the correlation functions along the axes perpendicular Δr_⊥_ and parallel Δr∥ to the preferred orientation revealed that, as the system evolved towards an aligned state, the discrepancy in the decay behavior of the correlation functions became more pronounced (Ext. Data Fig. 2d,e,g,h). This trend was observed in both experiments and simulations.

We extracted a length scale ξ from the correlation functions by determining the length when the correlation function reached the value 1/ϵ. In experiments, the correlation along both axes became more persistent over time (Ext. Data Fig. 2f). However, the perpendicular length xi_⊥_ grew more monotonically, while the parallel length xi∥ was strongly connected to the multiple phases. In contrast, in simulations, only the parallel axis showed a stronger correlation in the parallel direction, while the perpendicular axis remained unaffected (Ext. Data Fig. 2i). The cause for this is that as the monolayer matures, the phenomenological properties of the cell layer changed slightly over time, increasing the active length scale. This is not incorporated into the model; hence, the growth parallel set by the active length scale didn’t change.

### String visualization in simulations

The nematic director n = [cos(α), sin(α)] was defined both in orientation region region 0 ≤ α_1_ ≤ π (first and second quadrant) and −π/2 ≤ α_2_ ≤ π/2 (first and fourth quadrant). The parallel and perpendicular strings were defined to go between two resolution points if the absolute angle difference |Δα_1_| or |Δα_2_| was larger than π/2, respectively.

### Simulations

To complement the experiments, we simulated the cell layers as an active nematic film, where the third-dimensional shear flow was incorporated as an aligning field. We solved the continuum equations using a hybrid lattice Boltzmann approach [38]. We assumed that the active nematic film flows with collective velocity 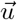 and has long-range orientational order described by the tensor order parameter ***Q***, which mimics the cell alignment.

The dynamics of the orientational order parameter ***Q*** at each position 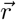 and time t are described by the Beris-Edwards transport equation

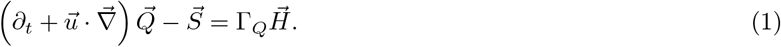

The co-rotation term 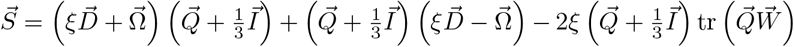 determines the align-ment of the cells in response to gradients in the velocity field, with **Ω** the rotational part and ***D*** is the extensional part of the velocity gradient tensor 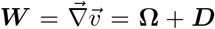. The alignment parameter ξ is considered deep in the flow aligning regime and set to ξ = 0.9 as cells align strongly with themselves.

The molecular field 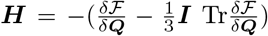 is a functional derivative of free energy density *ℱ*, describing the relaxation towards equilibrium at a rate Γ_*Q*_. The free energy consists of a Landau-De Gennes contribution, 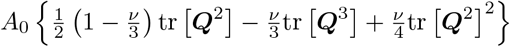 with A_0_ = 0.05, and Frank-Oseen deformation 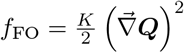 with K = 0.02. We used ν = 2.55, which favors the isotropic state in the absence of activity or any aligning field. Lastly, it contains a field strength f_Field_ = −ϵ_0_***E*** *·* ***Q*** *·****E*** where ***E*** is a matrix setting the direction of the flow and ϵ_0_ is the strength of the field that induces nematic ordering along the flow axis.

The system also obeys the Navier-Stokes equations for the velocity field within the active film. Assuming constant fluid mass density ρ (not active material concentration ϕ) leads to the incompressibility condition

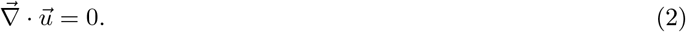

and

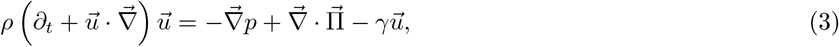

where p is the pressure and **Π** is the stress tensor which includes the standard viscous stress **Π**^visc^ = 2η***E*** for film viscosity η = 2/3. Furthermore, it contains the elastic stress due to the nematic nature of the order parameter

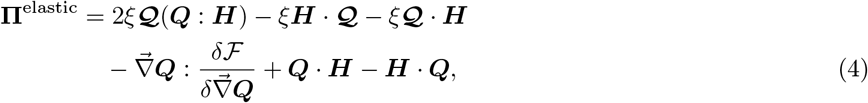

where ***𝒬*** = ***Q*** + ***I***/3. Lastly, the stress contains an active component

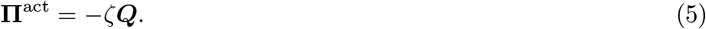

We used ζ = 0.03 to induce weak nematic ordering in the absence of an aligning field.

## II. LIST OF SUPPLEMENTARY MOVIES

**Supplementary Movie 1:** Phase contrast images of HAOECs exposed to 20 dyn/cm^2^ shear stress flowing from left to right. The HAOECs align in the direction of the flow.

**Supplementary Movie 2:** Phase contrast images of HAOECs from Supplementary Movie 1 with an verlay of the nematic director field and the topological defects as the cells reorient in the flow direction.

**Supplementary Movie 3:** Simulations of the nematic director field, colored by domain, were conducted with an anisotropic field strength of ν = 0.25 along the horizontal axis and an activity set to ζ = 0.03.

**Supplementary Movie 4:** Phase contrast images of HAOECs with an overlay of the domains. The defects are connected by red and green strings, indicating the energy intensive transition and the energy efficient transition between the domains.

**Supplementary Movie 5:** Simulations of the nematic director, colored by domain, from Supplementary Movie 3, with an overlay of defects that are connected by red and green strings indicating the energy intensive transition and the energy efficient transition between the domains.

## III. EXTENDED DATA

**Extended Data FIG. 1.**
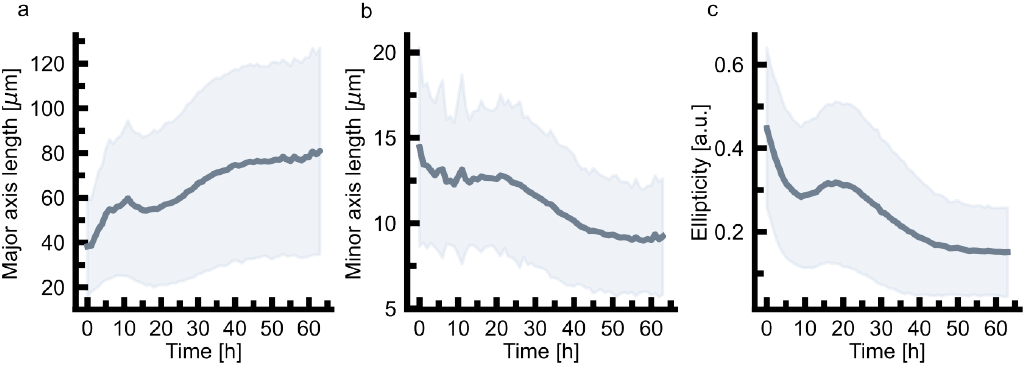
(a) The major axis length of HAOECs during exposure to shear flow. (b) The minor axis length of HAOECs during exposure to shear flow. (c) The ellipticity of HAOECs during exposure to shear flow. An ellipticity of 1 corresponds to a perfect circle.

**Extended Data FIG. 2.**
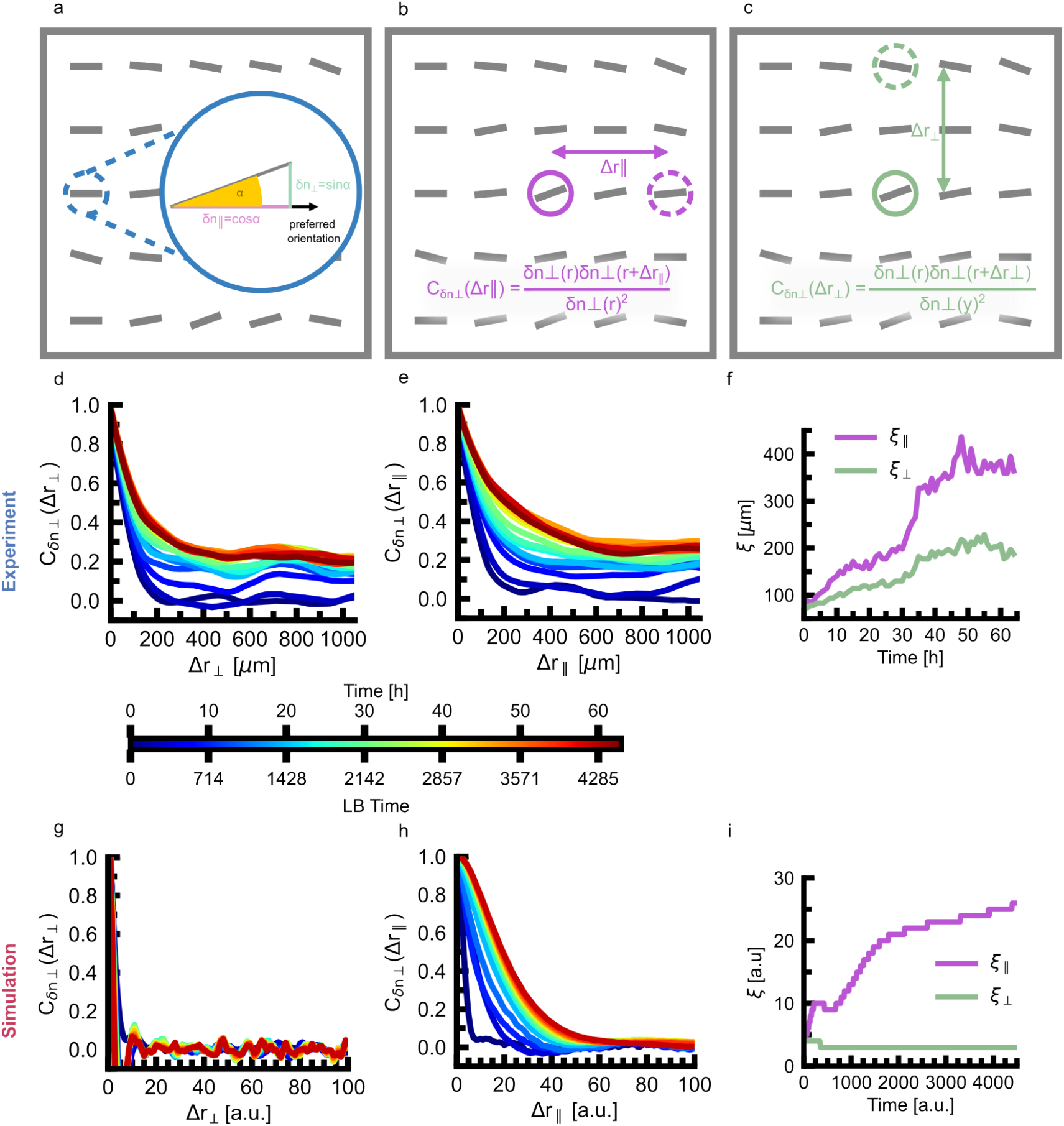
Correlation dynamics reveal anisotropic growth of similarly aligned regions. (a) Projections of the nematic director from the preferred axis of orientation 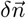. (b) Anisotropic growth of correlation length parallel Δr_∥_ and (c) perpendicular Δr_⊥_ to the preferred axis. (d) Time evolution of the perpendicular C_*δn*⊥_(r_⊥_) and (e) parallel C_*δn*⊥_(r_∥_) correlation function for the experimental data, (f) with corresponding correlation lengths. (g-i) Corresponding simulation figures to figures d-g.

**Extended Data FIG. 3.**
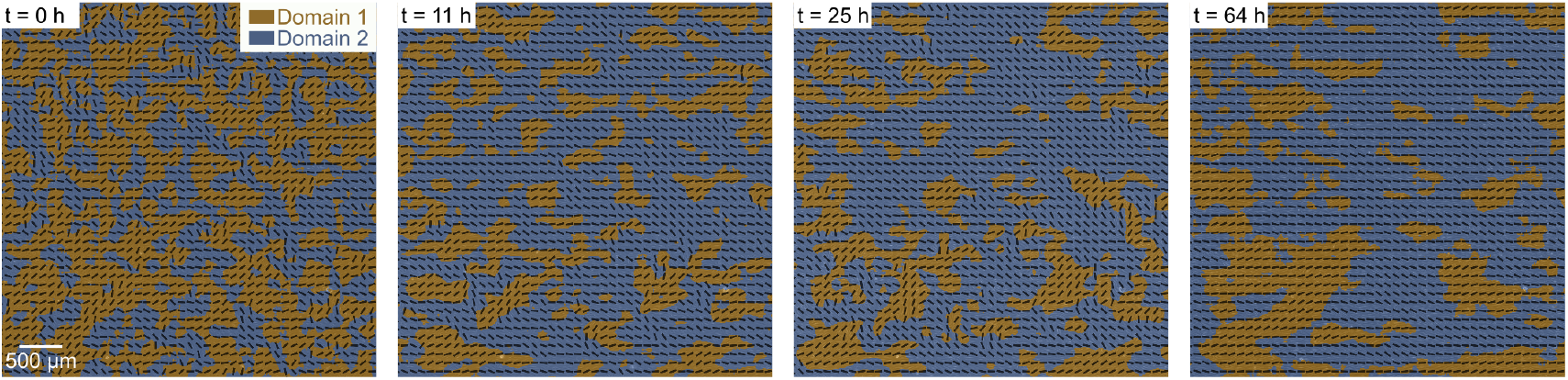
Coarsening dynamics of domains. Domains start symmetrically spaced, but over time, they coarsen anisotropically. The coarsening of the domains in the experimental data halts (t=11h to t=25) during the intermediate disordering phase (II) and progresses during the initial ordering phase (I) (t=0 to t=10h) and the steady-state ordering phase (III) (t=26 to t=64h), during which domains become more anisotropically shaped.

## Notes

### Competing Interest Statement

The authors have declared no competing interest.

